# ConfUSIus: an accessible and scalable toolkit for functional ultrasound imaging analysis

**DOI:** 10.64898/2026.10.27.762250

**Authors:** Samuel Le Meur-Diebolt, Felipe Cybis Pereira

## Abstract

Functional ultrasound imaging (fUSI) maps brainwide hemodynamics at high resolution, yet analysis requires expert knowledge as pipelines remain fragmented across laboratories. We introduce ConfUSIus, an open-source Python package and graphical interface for accessible, scalable, and reproducible fUSI analysis. Its common data model unifies heterogeneous fUSI recordings across beamformed IQ processing, motion correction, registration, denoising, and statistical analysis. ConfUSIus lowers barriers and facilitates method development, comparison, and sharing.

---

Functional ultrasound imaging (fUSI) has matured into a powerful tool for systems neuroscience, while opening new avenues for clinical translation. In animals, fUSI maps hemodynamics across large fields of view at high spatiotemporal resolution [1]. These capabilities have helped reveal how visual objects refine head-direction coding [2] and how olfactory cues engage reproductive circuits [3]. Combined with electrophysiology, fUSI has refined our understanding of neurovascular coupling [4], [5]. It has also revealed striking hemodynamics during sleep, from REM-related surges in rats to slow rhythms in mice and reptiles [6], [7]. In humans, studies have demonstrated bedside imaging of neonatal brain activity [8], cortical imaging through cranial windows [9] and brain monitoring during walking [10].

Yet the analysis infrastructure for fUSI lags behind advances in imaging technology. Recordings come in vendor-or laboratory-specific formats, and analysis often relies on custom pipelines that are difficult to maintain, reproduce and share. This fragmentation complicates anatomical alignment, multimodal synchronization, and comparisons across laboratories, which often depend on missing spatial and temporal metadata. Open tools such as PyfUS [11] and OfUSA [12] have made fUSI analysis more accessible, but cover only parts of the analysis workflow, leaving important steps such as beamformed IQ processing and automated atlas registration outside their scope. Therefore, researchers still lack a common, scalable framework linking heterogeneous recordings and analyses, from signal extraction to statistical inference.

To address this gap, we developed ConfUSIus, an open-source Python package and graphical interface for accessible, scalable and reproducible fUSI analysis (Fig. 1). ConfUSIus supports recordings from commercial imaging systems and custom laboratory setups, covering hemodynamic analysis from beamformed IQ processing to statistical modeling. Instead of a fixed pipeline, researchers can run its components independently or combine them into workflows tailored to different experimental designs.

**Fig. 1.**
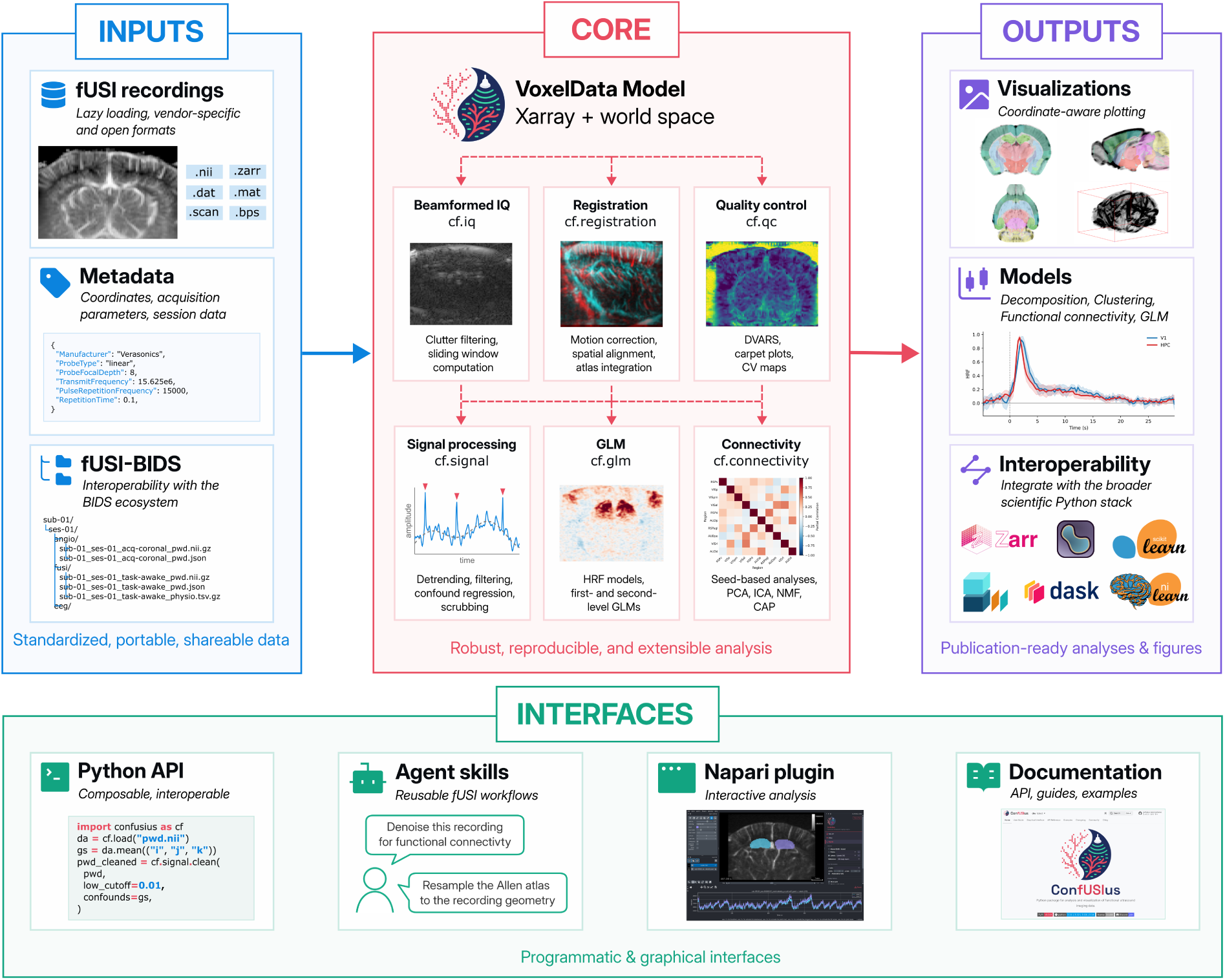
ConfUSIus framework overview. ConfUSIus unifies heterogeneous fUSI recordings and metadata in a common data model (VoxelData). Programmatic, agentic, and graphical interfaces provide scalable preprocessing, registration, quality control, statistical analysis, and visualization.

ConfUSIus connects ultrasound signal processing to the study of brain function (Fig. 1). From beamformed IQ recordings, researchers can recover complementary information about tissue and blood dynamics. Motion correction, denoising and registration help make these measurements reliable and comparable across time, sessions and anatomical regions. Researchers can then investigate stimulus-evoked responses, spontaneous brainwide activity and their links to behavior. These workflows can be scripted in Python, explored interactively in napari [13], or supported by reusable agent skills for AI-assisted analysis.

Underlying ConfUSIus is VoxelData, an Xarray-based data model that connects fUSI recordings and their derivatives throughout analysis (Fig. 1). From beamformed IQ and power Doppler recordings to anatomical atlases and statistical maps, VoxelData preserves physical coordinates, anatomical transformations and acquisition metadata, including the geometry and timing of multiple probe positions. Xarray keeps VoxelData interoperable with the broader PyData ecosystem, allowing researchers to use established Python tools without converting to a separate data structure [14]. To process large recordings efficiently, ConfUSIus uses Dask [15] for lazy, parallel and out-of-core computation on volumetric and beamformed IQ datasets reaching hundreds of gigabytes. VoxelData also supports the fUSI-BIDS specification [16], promoting consistent data organization and metadata to facilitate sharing and integration with complementary modalities such as electrophysiology and behavior [17].

We next describe how ConfUSIus addresses key challenges across fUSI processing and analysis, from extracting physiological signals to investigating brainwide hemodynamics. We illustrate these capabilities using published datasets from mice, rats and humans [5], [9], [18], [19] (Fig. 2).

**Fig. 2.**
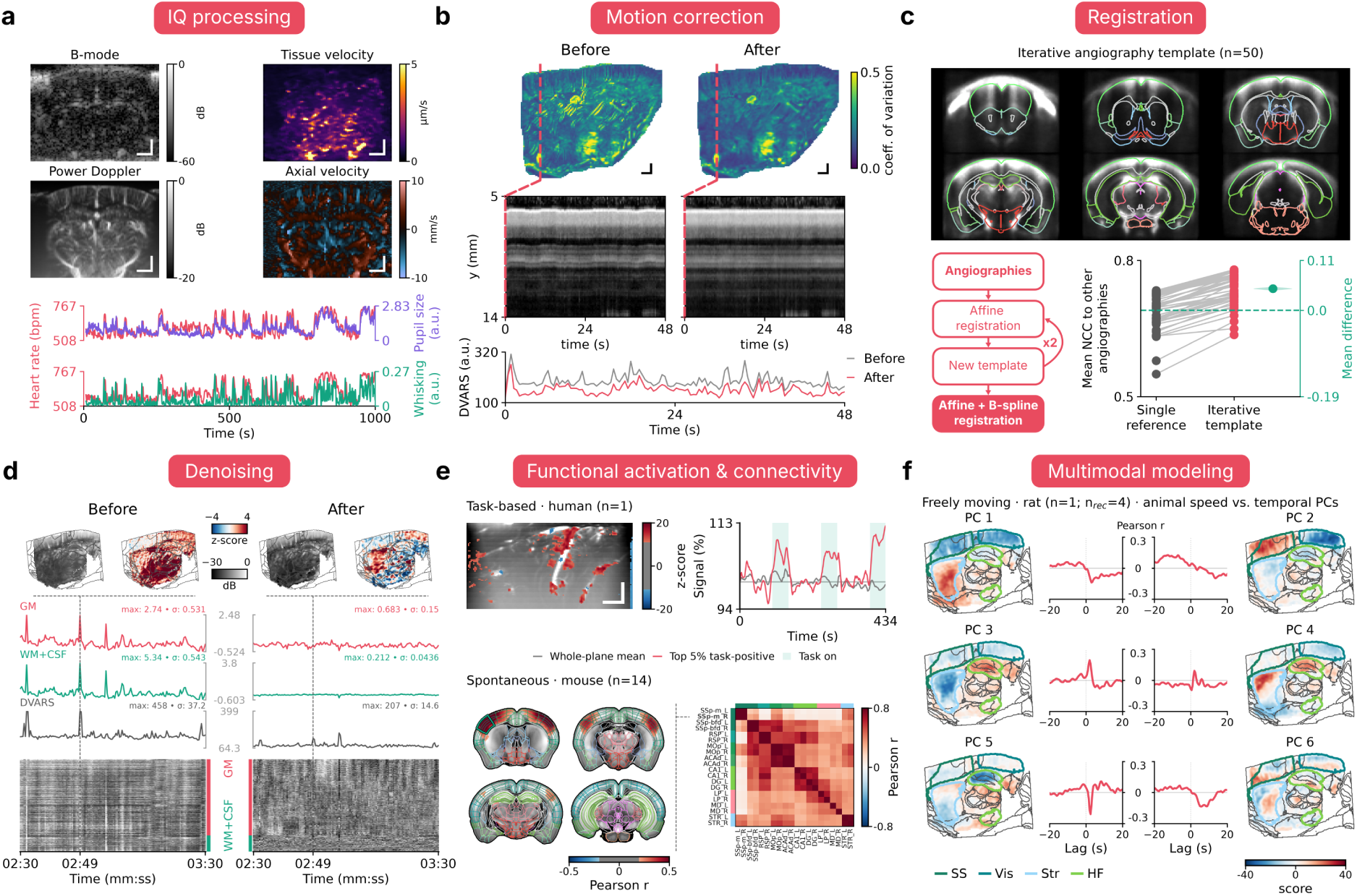
ConfUSIus applications across species and paradigms. **a**, Beamformed IQ processing yields B-mode anatomy, power Doppler, axial tissue and blood velocities, and physiological signals such as heart rate. IQ-derived heart rate is shown alongside simultaneously measured pupil size and whisking. **b**, Volume-wise motion correction reduces spatial and temporal variability. **c**, Vascular atlas registration using study-specific templates improves alignment consistency. Across 50 mouse angiographies, the iterative-template strategy increased mean inter-angiography correlation relative to single-reference registration (paired mean difference, 0.048; 95% bias-corrected and accelerated bootstrap confidence interval, 0.044–0.052; 5,000 paired resamples). **d**, Denoising reduces widespread fluctuations, assessed with z-score maps, gray matter (GM) and white matter/cerebrospinal fluid (WM+CSF) traces, DVARS and carpet plots. **e**, Task-evoked activation from general linear modeling in one human participant and spontaneous functional connectivity in 14 mice. **f**, Principal component analysis of four recordings from one freely moving rat reveals spatial patterns and their lagged cross-correlations with animal speed.

Beamformed IQ signals carry more than the blood-volume information retained in power Doppler (Fig. 2a). However, processing these high-volume-rate signals requires substantial computation and memory management. ConfUSIus uses Dask-backed, out-of-core functions to extract B-mode anatomy, power Doppler, and axial tissue and blood-flow velocities, from which physiological readouts such as heart rate can be estimated [20]. In our example, IQ-derived heart rate covaries with pupil size and whisking, illustrating how these readouts can be related to behavioral state (Fig. 2a). With standard defaults, users can adjust clutter filtering, integration, and sampling rates, or apply custom functions without implementing windowing or parallelization. Because clutter filtering can strongly affect signal quality and downstream analyses, this flexibility enables systematic comparisons of beamformed IQ processing methods [21], [22].

Beyond signal extraction, reliable fUSI analysis requires spatial alignment within and across recordings. Within recordings, brain motion can introduce spurious signal fluctuations. ConfUSIus uses parallelized volume-wise registration to correct those displacements and estimate motion, reducing artifactual variability (Fig. 2b). Across sessions and animals, registration is more challenging because vascular images differ in contrast and standardized vascular references remain limited [17]. For this, ConfUSIus provides automated registration methods and published vascular templates aligned with the Allen Mouse Brain Atlas [19], [23], [24]. Researchers can further refine alignments interactively through the graphical interface. Rather than aligning all animals to a single reference angiography, researchers can also build study-specific vascular templates by iteratively registering and aggregating angiographies [19], [22], [25]. In 50 mice, this approach improved registration metrics significantly (Fig. 2c).

Spatially aligned fUSI signals may still require denoising to reduce artifacts while preserving meaningful hemodynamic fluctuations. ConfUSIus provides configurable denoising workflows combining temporal filtering, motion scrubbing, and confound regression, as recommended by recent evaluations of fUSI denoising strategies [22]. Quality metrics and carpet plots help assess the effects of preprocessing parameters. In a recording from a freely moving rat [18], we show that denoising reduces widespread motion-related fluctuations and reveals more localized hemodynamic patterns in the striatum and visual cortex (Fig. 2d).

Finally, ConfUSIus brings together complementary approaches to studying brain function. Its statistical framework combines hypothesis-driven analyses, including general linear models and region-based functional connectivity, with unsupervised methods that identify distributed hemodynamic patterns. We illustrate these approaches by mapping task-evoked responses in a human participant and spontaneous functional networks in 14 mice (Fig. 2e). We also applied principal component analysis to 4 recordings from a freely moving rat, revealing spatial patterns whose time courses correlated with animal speed at different temporal lags (Fig. 2f).

Despite its broad capabilities, some challenges remain tied to experimental conditions and computational requirements. Anatomical registration requires suitable vascular templates, currently most developed for mice, and may need manual initialization. Optimal preprocessing choices also depend on acquisition conditions and scientific objectives. IQ processing currently targets offline parallel analysis rather than real-time applications such as ultrasound brain-machine interfaces [26], [27]. Future developments will expand reference datasets, support systematic comparisons across experimental settings and introduce GPU-accelerated processing.

Altogether, ConfUSIus provides a common foundation for accessible, scalable, and reproducible fUSI analysis. Its modular architecture makes it easier to develop and reuse analysis methods across laboratories. Beyond fUSI, VoxelData natively supports ultrasound localization microscopy (ULM) maps, paving the way for ULM to benefit from the same processing, visualization, and analysis tools [28], [29]. With open datasets, documented workflows, and community support, ConfUSIus can help move fUSI beyond laboratory-specific pipelines toward methods that can be systematically tested, compared, and trusted.

## Online methods

### VoxelData model

ConfUSIus uses the VoxelData model to represent spatially referenced recordings and their derivatives. Built upon Xarray [14], VoxelData is inspired by the SpatialData model from the scverse ecosystem [30]. Rather than introducing a new container format, VoxelData defines a set of conventions that any Xarray DataArray can follow, verifiable through dedicated validation functions. These conventions associate volumetric arrays with physical geometry, coordinates along additional dimensions (such as time, probe position or vector components), and metadata following the fUSI-BIDS specification [16]. The same model can therefore represent beamformed IQ and power Doppler recordings, anatomical references, displacement fields, statistical maps and other spatially referenced data used in fUSI analyses.

VoxelData follows a consistent dimension convention with three mandatory spatial dimensions (*i, j, k*) and optional temporal, probe-position and additional dimensions. Spatial coordinates are defined by a custom Xarray index that maps voxel positions to physical space using a voxel-to-world affine transformation, or one transformation per probe position for multi-pose acquisitions. This representation accommodates arbitrary imaging orientations without requiring resampling. VoxelData can also store affine transformations linking its world coordinates to other reference spaces, such as laboratory coordinates or the Allen Mouse Brain Common Coordinate Framework. These transformations allow users to switch between coordinate systems without resampling the underlying data.

Temporal coordinates preserve acquisition timestamps, window durations and timestamp reference points. Accurate tracking of this information is particularly important for multimodal experiments, where fUSI recordings must be synchronized with external measurements such as electrophysiology, physiological signals or behavior. For sequential multi-pose acquisitions, VoxelData also supports posedependent timestamps and slice-specific timing information, preserving acquisition-time offsets between different brain locations.

Because VoxelData remains a standard Xarray DataArray, it integrates directly with the broader PyData ecosystem, allowing users to apply Xarray operations and pass arrays to compatible scientific Python libraries without conversion to a proprietary data structure. A dedicated ‘fusi’ Xarray accessor provides convenient access to spatial and temporal spacing, volume origin, orientation and fUSI-specific processing methods. Xarray also supports alternative computational backends, including GPU-based arrays such as CuPy for compatible operations. ConfUSIus leverages this flexibility through Dask [15], enabling lazy, chunked and out-of-core computation on large recordings.

### Data loading and standardization

ConfUSIus provides readers and conversion utilities for vendor-specific file formats from Iconeus, AUTC and EchoFrame-based systems. Recordings are converted into VoxelData arrays, preserving available imaging geometry, timing information and acquisition metadata. Other formats are supported by constructing VoxelData arrays directly from raw arrays using convenient helper functions.

For storage and exchange, ConfUSIus supports NIfTI and Zarr. NIfTI provides compatibility with established neuroimaging software, while Zarr offers chunked storage suited to large recordings, particularly beamformed IQ data. ConfUSIus also supports the fUSI-BIDS specification [16], providing utilities for standardized dataset organization, metadata validation, and integration of experimental events and physiological recordings.

### Datasets and anatomical references

All analyses used previously published fUSI recordings, converted to the fUSI-BIDS specification when necessary. Anatomical registration and template construction used angiographies from 50 mice, while functional connectivity analyses used sedated recordings from 14 mice [19]. Motion correction, denoising and principal component analysis were illustrated using four open-field recordings from one freely moving rat [18]. General linear modeling was demonstrated using one human recording acquired through an acoustically transparent cranial window [9], and beamformed IQ processing using an awake head-fixed mouse recording with simultaneous pupil and whisking measurements [5].

Anatomical references included the Allen Mouse Brain Common Coordinate Framework [24] and Waxholm Space rat atlas [31], accessed through the ConfUSIus interface to BrainGlobe [32], alongside published mouse vascular templates [19], [23].

To facilitate reproducibility and reuse, ConfUSIus provides a collection of six published fUSI datasets and two vascular templates hosted on the ConfUSIus Functional Ultrasound Imaging Dataset Collection (https://registry.opendata.aws/confusius/). Datasets are versioned and accompanied by documented provenance, conversion recipes and licenses. Built-in functions enable selective downloading, integrity verification and local caching for offline reuse.

### Beamformed IQ processing

ConfUSIus processes beamformed IQ recordings using configurable sliding temporal windows, with Dask providing parallel and out-of-core computation. Power Doppler and axial velocity estimation use nested windows, allowing clutter filtering and signal estimation to be controlled independently. Available clutter filters include fixed-rank and adaptive singular value decomposition (SVD), as well as temporal Butterworth filtering. B-mode intensity is computed from the mean IQ magnitude, power Doppler from the mean squared magnitude of clutter-filtered IQ signals, and axial velocity from the phase of the temporal autocorrelation of complex IQ signals [33]. Importantly, users can also define custom processing functions operating on a single block of IQ data, while ConfUSIus automatically handles windowing, parallelization, and processing across the entire recording.

For Fig. 2a, we processed beamformed IQ data from an awake, head-fixed mouse [5]. B-mode, axial tissue and blood velocity, and power Doppler signals were averaged over the first 30,000 beamformed IQ frames, removing the 40 strongest SVD components for blood measurements. To illustrate the extraction of additional physiological information, we estimated the cardiac rhythm following the PhysiofUS procedure [20]. Briefly, we identified a cardiac-related spatial component by decomposing axial velocity fluctuations over an initial 4-second window. The corresponding spatial mask was used to extract a velocity time course from the recording. Heart rate was estimated from autocorrelation peaks within the 8–14 Hz frequency band using 3-second windows with a 0.25-second stride. The resulting estimates were smoothed and temporally aligned with simultaneously recorded pupil size and whisking measurements.

### Anatomical registration

ConfUSIus provides a general-purpose image registration framework built on SimpleITK [34], supporting alignment across recordings, animals and anatomical reference spaces. It supports translation, rigid, affine and B-spline transformations, with normalized cross-correlation (default) or Mattes mutual information as similarity metrics. Registration can be restricted to selected regions using masks and initialized from image centers, centers of mass or user-defined transformations. Manual initializations can be performed using the ConfUSIus GUI. The framework returns the registered image, estimated transformation and convergence diagnostics. Transformations can subsequently be applied to other volumes, independently of their spatial grids, facilitating the alignment of anatomical annotations, masks, and functional maps.

The published mouse vascular templates [19], [23] provided within ConfUSIus store transformations to the Allen Mouse Brain Common Coordinate Framework [24]. Registering a recording to one of these templates therefore aligns it with the atlas, allowing anatomical annotations and reference images to be mapped into recording space.

For Fig. 2c, we constructed a study-specific vascular template from 50 mouse angiographies [19]. Each angiography was divided by its mean intensity and raised to the power 0.25 to reduce the influence of high-intensity vessels. Starting from the voxel-wise median of the unregistered angiographies, we performed three iterations of affine registration to the current template, followed by voxel-wise median aggregation. The resulting template was symmetrized by registration to its mirror image and averaging. We compared this approach with registration to a single reference angiography, using normalized cross-correlation and an additional B-spline registration with a 5 × 5 × 5 control-point mesh for both strategies. Registration consistency was quantified by the mean normalized cross-correlation (NCC) of each registered angiography with all others, within their common field of view and an Allen atlas-derived brain mask. Differences between strategies were evaluated using a paired bias-corrected and accelerated bootstrap with 5,000 resamples.

For Fig. 2b and Fig. 2d, we used the ConfUSIus napari plugin to draw a brain mask and manually align the recording to the Waxholm Space rat atlas [31].

### Volumewise motion correction

ConfUSIus uses the same registration framework to correct brain motion within recordings by independently aligning each volume to a common reference, selected from the recording or provided separately. Individual registrations are parallelized across CPU cores using joblib [35]. The corrected recording retains its original spatial grid and metadata, while estimated transformations and convergence diagnostics are returned separately. Motion parameters can also be used to calculate framewise displacement for quality control, motion scrubbing and confound regression [36].

For Fig. 2b, we registered 120 consecutive volumes from the freely moving rat recording to its first volume, using rigid transformations, normalized cross-correlation and an optimizer step size of 1. Motion correction was evaluated within a brain mask using voxel-wise coefficients of variation and DVARS, as described under Quality control.

### Quality control

ConfUSIus provides spatial and temporal quality-control metrics to identify artifacts and assess preprocessing outcomes. DVARS quantifies signal changes between consecutive volumes using the root-meansquare intensity difference across brain voxels [36], with optional standardization accounting for temporal autocorrelation [37]. Spatial variability is characterized by the voxel-wise coefficient of variation (CV), defined as the temporal standard deviation divided by the temporal mean. Although temporal signal-to-noise ratio (tSNR) is also available, its interpretation can be misleading for power Doppler recordings, particularly in regions with low vascular signal [22]. Carpet plots provide complementary visualization of voxel-wise time series, revealing widespread signal fluctuations and transient artifacts.

For Fig. 2b, we compared CV maps and DVARS before and after motion correction within a brain mask. For Fig. 2d, we assessed the effects of denoising using DVARS, gray matter and white matter/ cerebrospinal fluid signal traces, and carpet plots.

### Signal denoising

ConfUSIus provides a flexible signal processing pipeline adapted from Nilearn, combining detrending, temporal filtering, motion scrubbing, confound regression and standardization [38]. This pipeline follows recommendations by Lindquist and colleagues to minimize the reintroduction of previously removed nuisance signals [39]. When scrubbing is enabled, censored volumes are temporarily interpolated before filtering and removed afterward. Confound regressors undergo the same detrending, filtering and censoring operations as the imaging signals before regression. The framework supports both predefined and data-driven confounds, allowing denoising strategies to be tailored to experimental conditions and to incorporate recent recommendations for mitigating motion artifacts in fUSI [22].

For Fig. 2d, we analyzed one of the four rat recordings. Following volume-wise motion correction, we extracted gray matter and white matter/cerebrospinal fluid masks from the Waxholm Space rat atlas [31], restricted to a manually defined brain mask. We applied high-pass filtering at 0.01 Hz and regressed the first 12 anatomical CompCor components estimated from white matter and cerebrospinal fluid [40], followed by spatial smoothing with a full-width at half-maximum of 0.2 mm. The effects of preprocessing were assessed using voxel-wise z-scored maps, tissue-averaged time courses, DVARS and carpet plots, highlighting the suppression of widespread motion-related fluctuations and the emergence of more localized hemodynamic patterns.

### General linear modeling

ConfUSIus implements voxel-wise first- and second-level general linear models adapted from Nilearn [38]. Design matrices combine experimental events or continuous regressors with nuisance confounds and temporal drift terms. Supported hemodynamic response functions include the Glover and SPM models, finite impulse response models and functions derived from published fUSI studies [41], [42]. Models support ordinary least squares or autoregressive noise estimation, fixed-effects combination across runs and group-level inference. Statistical contrasts return effect, variance and statistical maps as VoxelData arrays, with utilities for thresholding and multiple-comparison correction.

For Fig. 2e, we analyzed a human task recording [9], retaining depths of 10 to 30 mm to select visible brain regions. After temporal resampling and motion correction, we followed the original study’s preprocessing pipeline by applying Gaussian smoothing (0.47 mm full-width at half-maximum) and a six-volume moving average. Signals were normalized to the mean resting-state intensity. The model used a gamma hemodynamic response function (onset, 3 s; peak delay, 1.4 s; dispersion, 0.7 s), cosine drift regressors (0.002 Hz cutoff) and ordinary least squares.

### Functional connectivity

ConfUSIus supports seed-based connectivity maps and region-to-region connectivity matrices, with methods adapted from Nilearn. Seed-based maps compute Pearson correlations between voxel-wise signals and time series extracted from atlas-defined or user-defined regions, or supplied externally. Connectivity matrices support covariance, correlation, partial correlation, precision, and tangent-space estimates. These analyses can be combined with the denoising methods described above.

For Fig. 2e, we analyzed recordings from 14 sedated mice, each comprising four coronal planes sampled every 2.4 s [19]. After in-plane motion correction and temporal alignment, we registered recordings to the Allen atlas through study-specific functional and vascular templates. Signals were high-pass filtered at 0.01 Hz and low-pass filtered at 0.1 Hz, with volumes censored when framewise displacement exceeded 0.02 mm or relative DVARS exceeded 3 in any plane. We computed Pearson correlations between ten bilateral Allen regions and seed-based maps from the right primary somatosensory mouth area. Group results were averaged using Fisher’s z transformation, displaying seed maps only where at least half of the mice contributed data.

### Dimensionality reduction

ConfUSIus implements principal component analysis (PCA), independent component analysis (ICA) and nonnegative matrix factorization (NMF), adapted from scikit-learn for VoxelData arrays [43]. These methods support spatial and temporal decompositions, returning component maps with their original spatial coordinates and associated time courses. Users can restrict decomposition to selected brain regions and reconstruct recordings from the estimated components.

For Fig. 2f, we analyzed four recordings from a single freely moving rat. Following motion correction, recordings were aligned across sessions using iterative affine and B-spline registration and manually aligned to the Waxholm Space rat atlas [31]. Each recording was high-pass filtered at 0.01 Hz, denoised using 12 white matter/cerebrospinal fluid confound components, standardized and spatially smoothed (0.2 mm full-width at half-maximum). The recordings were then concatenated for spatial PCA, retaining six components. Animal body speed was derived from the tracking data released with the original dataset and smoothed over 1 s. We computed Pearson correlations between body speed and component time courses at lags of up to ±20 s, pooling results after within-session standardization.

### Software interfaces

ConfUSIus provides a Python API and an interactive graphical interface built as a napari plugin [13]. The Python API exposes processing and analysis functions directly or through the ‘fusi’ Xarray accessor, alongside Matplotlib-based utilities for static visualization. The napari plugin enables interactive exploration of spatial and temporal recordings, synchronized video playback, region-of-interest annotation, signal extraction, event annotation and quality control. Dedicated panels also support within-recording motion correction and registration between acquisitions, including visualization and reuse of estimated transformations. Interactive plots are linked to the image viewer, allowing users to navigate directly between signal traces and their corresponding imaging frames.

## Data availability

Publicly redistributable fUSI datasets and vascular templates are available through the ConfUSIus Functional Ultrasound Imaging Dataset Collection, hosted on Amazon S3 with support from the AWS Open Data Sponsorship Program (https://registry.opendata.aws/confusius/). Where possible, datasets have been organized according to the fUSI-BIDS specification and can be downloaded directly or accessed programmatically through ConfUSIus. Dataset preparation scripts, provenance information and licensing details are available at https://github.com/confusius-tools/confusius-datasets. Original data sources and associated publications are acknowledged in the collection documentation (https://confusius.tools/latest/user-guide/datasets/).

## Code availability

ConfUSIus is freely available as open-source software under the BSD 3-Clause license at https://github.com/confusius-tools/confusius and distributed through PyPI (https://pypi.org/project/confusius/). Software releases are archived on Zenodo (https://doi.org/10.5281/zenodo.18611124). Comprehensive documentation, installation instructions, API references and executable analysis examples are available at https://confusius.tools/. Code to reproduce the analyses and figures presented in this study is available at https://github.com/confusius-tools/confusius-paper.

## Acknowledgments

This work was funded by a Wellcome Trust Investigator Award to M. Carandini and K. D. Harris (grant 223144/Z/21/Z). We thank M. Carandini and A. Landemard for helpful suggestions. We thank the AWS Open Data Sponsorship Program for supporting the storage and distribution of the ConfUSIus Functional Ultrasound Imaging Dataset Collection. We thank M. S. Blumberg and B. Dwyer for feedback on the ConfUSIus GUI. We thank M. S. Blumberg, B. Dwyer, C. I. J. Ferreira, J. P. P. Barbosa and M. A. Romano-Silva for providing example Iconeus SCANv2 files that guided the development of the experimental loader.

## Author contributions

S.L.M.-D. conceived the project. S.L.M.-D. and F.C.P. jointly designed the software architecture, defined its scientific scope, developed the software and methodology, curated and analyzed datasets, performed validation, prepared figures, and wrote and revised the manuscript. Both authors contributed equally to this work.

